# Introduction of CG methylation in *E. coli* induces mutagenesis at AT base pairs

**DOI:** 10.64898/2026.01.24.701492

**Authors:** K Hains, A. Klimczyk, P Sarkies

## Abstract

In many eukaryotic species, DNA is methylated at the 5 position of cytosine to form 5mC, predominantly within CG dinucleotides. Despite being conserved since the dawn of eukaryotic life, 5mC is often lost from individual lineages, suggesting that it may have detrimental effects. One such effect is genotoxicity, through the effect of 5mC on the process of cytosine deamination and its repair. Additionally, enzymes that introduce 5mC (DNA methyltransferases, DNMTs) can also damage DNA through alkylation and oxidative stress, but how these genotoxic effects combine to influence mutagenesis is unclear. To investigate how mutagenesis changes upon methylation of CG dinucleotides we introduced high levels of CG methylation into the bacteria *E. coli*. 5mC induction increased mutation at CG dinucleotides consistent with increased C to T mutations. We also discovered that 5mC induction led to increased mutations at AT base pairs, specifically in the absence of the alkylation repair enzyme AlkB. This effect was specific to certain *E. coli* strains and was not dependent on the DNA repair enzyme RecA, so its exact mechanism remains unclear. Together, our work highlights multiple mutagenic consequences of DNMT expression, which might act as selective pressures for organisms to lose 5mC across evolution.

## Introduction

Methylation of cytosine at the 5^th^ position (5mC) is an ancient modification conserved across eukaryotes (de Mendoza *et al*, 2020). Most commonly, 5mC is found within the CG sequence context and this is likely ancestral to eukaryotes (Sarkies, 2022b). This enables it to be propagated during DNA methylation due to the activity of maintenance methyltransferases, that can recognise hemi-methylated DNA and introduce methylation onto the newly synthesised strand (Law & Jacobsen, 2010). As a result, 5mC can act as an epigenetic mark, which can be stably propagated through mitosis (Holliday *et al*, 1987). In mammals, 5mC within promoters is associated with gene silencing, and 5mC can also act to silence repetitive elements (Weber *et al*, 2007). However, across eukaryotes more generally, promoter methylation is not common. Indeed, the genome-wide distribution of 5mC evolves rapidly (Feng *et al*, 2010; Zemach *et al*, 2010; de Mendoza *et al*, 2020).

The broad distribution of 5mC across eukaryotes implies its importance in cellular function and indeed, complete loss of cytosine methylation is lethal in a range of different organisms (Schulz *et al*, 2018; Bewick *et al*, 2019b; Okano *et al*, 1999; Liao *et al*, 2015). Nevertheless, the presence of 5mC is not universal. The levels of 5mC can vary enormously even within a single phylum (Lewis *et al*, 2020; Bewick *et al*, 2017, 2019a). Moreover, both 5mC and the DNA methyltransferases that introduce it have been lost completely in a number of lineages across eukaryotes, with prominent examples of species lacking 5mC altogether including the yeasts *S. cerevisiae* and *S. pombe*, the nematode *C. elegans* and the fruitfly *Drosophila melanogaster* (Ponger & Li, 2005). In all of these cases the loss of 5mC has occurred independently, implying that there must be selective advantages to losing 5mC under some circumstances.

A key property of DNA methyltransferase activity that may contribute to the loss of 5mC is its propensity to threaten genome stability. The most straightforward of these is due to deamination of cytosine. Cytosine deamination to uracil is amongst the most common spontaneous damage that occurs to DNA in cells (Lindahl, 1996). As a result, highly conserved enzymes (UNG; uracil deglycosylases) exist that counteract this by removing uracil and triggering restoration of the original cytosine by base excision repair (Liu *et al*, 2002). However, 5mC deaminates at a faster rates than unmodified cytosine. Moreover, deamination of cytosine forms thymine, and processing of the resulting TG mismatch by the glycosylase TDG is substantially less efficient than uracil excision (Sarkies, 2022a). Methylated cytosine also poses other problems for genome stability, because it is more likely to lead to miscoding by the replicative DNA polymerases, particularly the leading strand polymerase Pol epsilon (Tomkova *et al*, 2018, 2024). As a result, methylated cytosine is mutagenic. Indeed, the pre-eminent source of somatic mutations, for example in cancer genomes, is C-T mutations in the CG context, which can be attributed to deamination of 5mC (Alexandrov *et al*, 2020). Moreover, over evolutionary timescales in mammals, CG base pairs are unstable, resulting in their underrepresentation across most genomic regions (Illingworth & Bird, 2009).

In addition to these mechanisms of mutagenesis, formation of 5mC has also been shown to cause DNA damage through other mechanisms. DNA methyltransferases (DNMTs) that form 5mC have the propensity to introduce alkylation damage at low levels, through off-target catalytic activity that introduces 3mC instead (Rošić *et al*, 2018; Dukatz *et al*, 2019). This modification interferes with base pairing and is therefore a highly toxic form of DNA damage (Sedgwick, 2004). As a result, the occurrence of cytosine DNMTs across species correlates with the presence of the enzyme ALKB2, a specialised repair enzyme that directly oxidises 3mC to reform cytosine (Rošić *et al*, 2018; Lewis *et al*, 2020). Loss of ALKB2 when DNMTs are active leads to accumulation of 3mC (Rošić *et al*, 2018).

Recently, we developed a system to test the genotoxic effects of DNMT activity by introducing CG methylation into *E. coli*, to enable us to study the consequences for sensitivity to a range of different DNA damaging agents. To do this we introduced the prokaryotic methyltransferase M.SssI into *E. coli* compromised for its restriction modification system to allow stable propagation of *E. coli* with genome-wide CG methylation (Krwawicz *et al*, 2025). As predicted from the ability of DNMTs to introduce 3mC, expression o*f* M.SssI led to hypersensitivity to alkylation damage by methane methanesulfonate (MMS). Surprisingly, it also sensitized cells to oxidative stress, due to the fact that reactive oxygen species react with 5mC to produce 5fC, which is recognised as DNA damage by *E. coli* (Krwawicz *et al*, 2025). These data confirm that DNMTs have severe consequences for genome stability.

The propensity of DNMTs to damage DNA directly suggests that in addition to the well characterised mutagenicity of the 5mC base itself through deamination and miscoding, there may be other forms of mutagenesis due to DNMT activity. Unrepaired 3mC has been shown to lead to mutations, probably due to translesion DNA polymerases that promote error-prone bypass (Sikora *et al*, 2010). Moreover, unrepaired 5fC or other oxidative products of 5mC could result in mutations. We therefore set out to investigate how the mutation rate is affected by expression of M.SssI in *E. coli* and, using a sequencing-based assay, determine how the spectrum of mutations is affected. By combining expression of M.SssI with mutation of the DNA repair enzyme AlkB (homologous to eukaryotic ALKB2), we uncover an unexpected form of DNA mutations dependent on M.SssI expression that only appears in *ΔalkB* mutants. Intriguingly, this appears at AT bases. Our work indicates that the ability of DNMTs to damage DNA has more widespread mutagenic consequences than directly due to processing of methylated cytosine, potentially contributing to its frequent loss across evolution.

## Results

### Constitutive induction *of* M.SssI introduces mutations in *E. coli*

To investigate mutagenesis due to M.SssI expression we used a model that we developed previously, where the prokaryotic CG dinucleotide-specific DNMT M.SssI is expressed on a plasmid in *E. coli* cells derived from the C2523 cloning strain (Figure 1A-C). These cells lack the *mcrABC* restriction modification system, enabling stable maintenance of the DNMT-containing plasmid (Krwawicz *et al*, 2025). To investigate mutagenesis, we used a modified rifampicin resistance assay (Figure 1D; see methods). We grew *E. coli* expressing the M.SssI plasmid and then plated out onto plates containing rifampicin. Point mutations in *rpoB* can result in resistance to rifampicin. We assayed mutation rate as the fraction of resistant colonies. Expression of *m*.*sssI* led to an increase in the mutation rate (Figure 2A). To test if alkylation damage induction by DNMTs could result in mutations, we introduced M.SssI into cells where the AlkB DNA repair enzyme that repairs DNA alkylation damage had been deleted (*ΔalkB*). Induction of M.SssI led to a greater increase in mutation rate in the *ΔalkB* cells than in WT C2523 (two-way anova for interaction between M.SssI and *ΔalkB* p<1e-3). This implies that there are additional mutations that occur due to M.SssI activity when *alkB* is inactive.

**Figure 1.**
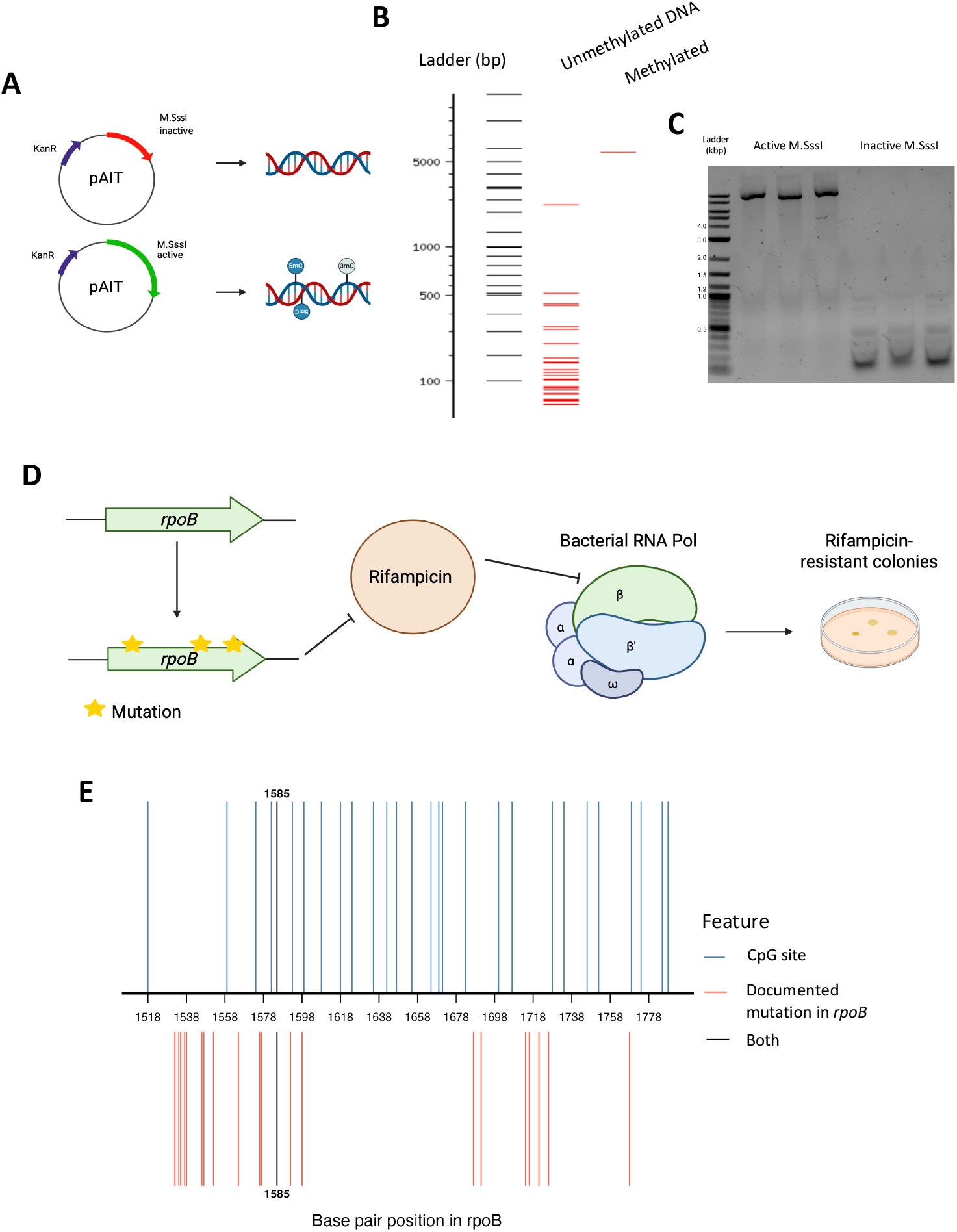
Constitutive CpG methyltransferase system and mutation selection assay in *E. coli*. A) Plasmids containing either full length active or truncated inactive versions of the methyltransferase M.SssI (pM.SssI) for transformation into *E. coli*, introducing genome-wide CpG methylation. Created in Biorender. B) Schematic of methylation-sensitive HpaII restriction enzyme digest of plasmid DNA from *E. coli* containing either active or inactive M.SssI. HpaII is blocked by methylation in CCGG context. C) Plasmid DNA was isolated and digested by HpaII, then run on a 1% agarose gel. Digestion was blocked by CpG methylation in DNA from *E. coli* with active M.SssI, resulting in a higher band pattern compared to unmethylated, digested DNA. D) Schematic of rifampicin resistance assay workflow. Resistance to the antibiotic rifampicin was used to select for mutant *E. coli*. Off-target activity of M.SssI can result in mutagenic lesions such as 3mC. Mutations in *rpoB* alter the structure of the β subunit of RNA Polymerase and prevent the binding of rifampicin. This renders *E. coli* resistant to rifampicin and allows for selection of mutant colonies on rifampicin media. Created in Biorender. E) Documented mutational hotspots in *rpoB* against CpG sites by base pair. Overlapping sites shown in black.

**Figure 2.**
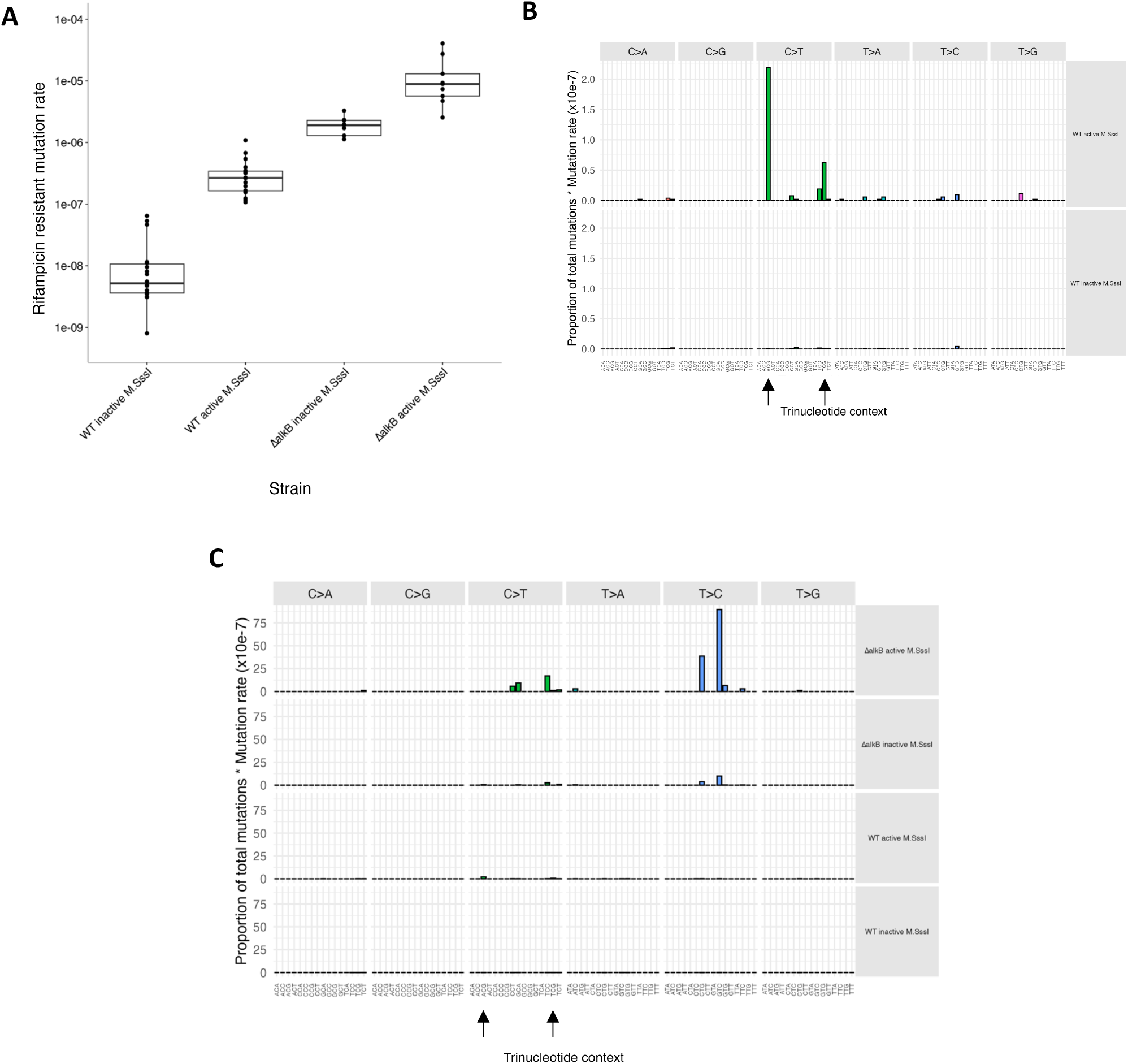
Introducing CpG methylation into C2523 *E. coli* increased mutation rate and resulted in a distinct non-cytosine mutational signature dependent on loss of *alkB*. A) WT and Δ*alkB* C2523 *E. coli* with active and inactive M.SssI were plated on both rifampicin and LB media. The number of rifampicin-resistant mutant colonies was compared to total colonies on LB media to calculate a rifampicin-resistant mutation rate. Each individual point represents an independent biological replicate, each averaged over two technical replicates. DNA from mutant colonies was isolated and a 500bp region of *rpoB* amplified by PCR. PCR products were sequenced to determine mutational signatures in WT (B) and WT and Δ*alkB* (C) C2523 *E. coli* with active and inactive M.SssI. Signatures are categorised by substitution type and trinucleotide context, comprised of the 3’ and 5’ bases adjacent to the mutated base. This gives 96 possible signatures. The percentage of unique signature was multiplied by average mutation rate for each strain to give a qualitative contribution to overall mutation signature. Observed signatures occurring in the CpG context are indicated by black vertical arrows.

### Mutations at AT base pairs occur due to M.SssI activity in cells lacking AlkB activity

To investigate what these mutations were, we sequenced a region of *rpoB* in resistant clones and tabulated the frequencies of different types of mutations. Importantly, a restricted number of point mutations are able to give rise to rifampicin resistance, so this does not represent an unbiased spectrum of mutations (Figure 1E). However, by comparing the spectrum of mutations that occur in WT in the presence or absence of M.SssI, we could infer the effect of M.SssI expression. M.SssI expression led to a large increase in the mutation rate at CG dinucleotides, consistent with the known effect of 5mC in increasing CG->TA mutations and the sequence specificity of M.SssI for CG dinucleotides. No marked increases in other mutation types were observed (Figure 2B).

In *ΔalkB* cells, the mutation spectrum was altered slightly compared to WT in the absence of *m*.*sssI*. However, when *m*.*sssI* was expressed there was a notable increase in mutations at AT base pairs, specifically AT-GC mutations, implying that M.SssI was responsible for AT mutagenesis but only in cells lacking *alkB* (Figure 2C). This was surprising because previously alkylation damage was shown to occur at cytosine in the form of 3mC, which would be expected to alter mutation rate at CG base pairs.

### *E. coli* strain background affects the mutational spectrum

As the induction of mutations at AT base pairs by M.SssI was surprising, we set out to investigate it in another context. We previously developed a system to induce M.SssI under the control of an arabinose-sensitive promoter on the pBad expression system (Krwawicz *et al*, 2025). This system cannot be used in the C2523 strain, so we introduced it into WT Top10 cells (Figure 3A, B). Induction of M.SssI for two hours led to an increase in mutation rate compared to uninduced cells grown in glucose (Figure 3C). We then engineered the *ΔalkB* mutation into Top10. This led to a further induction in mutation rate that was greater than the combined effect of *ΔalkB and* M.SssI alone (three-way anova for interaction between M.SssI, *ΔalkB* and arabinose p < 1e-4) (Figure 3C). Thus, M.SssI induced mutations that were dependent on loss of *alkB*, just as in the constitutive expression of M.SssI. However, when we sequenced rifampicin resistant colonies, we observed a large increase in CG mutations in the *ΔalkB* background but no evidence of mutations in the AT context (Figure 3D, E).

**Figure 3.**
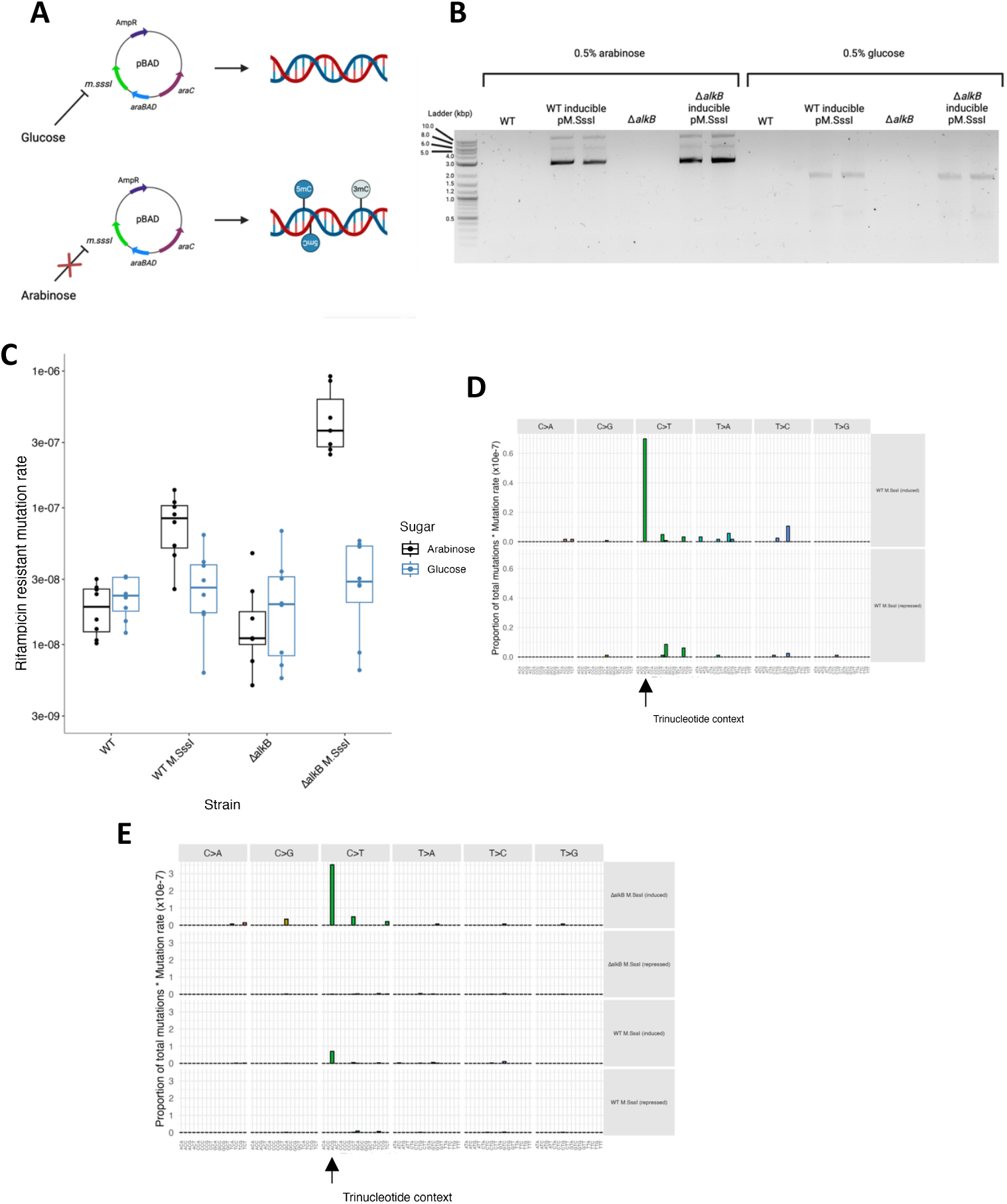
Inducing CpG methylation in Top10 *E. coli* increased mutation rate but did not introduce previously observed non-cytosine mutational signature. A) Schematic of engineered plasmid containing the full length, active M.SssI under the control of an arabinose-inducible P_BAD_ promoter (inducible pM.SssI). The promoter is repressed and M.SssI expression inhibited in the presence of glucose. Removal of glucose and addition of arabinose alleviates repression and induces M.SssI expression, leading to DNA methylation. Created in Biorender. B) HpaII restriction enzyme digest of plasmid DNA extracted from Top10 *E. coli* containing inducible M.SssI under 0.5% glucose or arabinose respectively. C) Rifampicin-resistant mutation rates were calculated for WT and Δ*alkB* Top10 *E. coli* transformed with inducible pM.SssI and treated with either 0.5% arabinose or glucose. Mutational signatures for WT (D) and WT and Δ*alkB* (E) with inducible pM.SssI treated with either 0.5% arabinose (induced) or glucose (repressed). Signatures are categorised by substitution type and trinucleotide context, comprised of the 3’ and 5’ bases adjacent to the mutated base. This gives 96 possible signatures. The percentage of unique signature was multiplied by average mutation rate for each strain to give a qualitative contribution to overall mutation signature. Observed signatures occurring in the CpG context are indicated by black vertical arrows.

One possible explanation for this discrepancy could be that the induction system led to different expression of M.SssI, affecting the mutational profile. However, when we introduced the constitutively expressed M.SssI into Top10 using the same plasmid as for the C2523, we observed increased mutation rate (Figure 4A) but no evidence of DNMT-dependent mutations at AT base pairs in *ΔalkB* mutant cells (Figure 4B). We therefore concluded that the mutations at AT base pairs due to DNMT activity depend on the cell background as well as the presence of AlkB.

**Figure 4.**
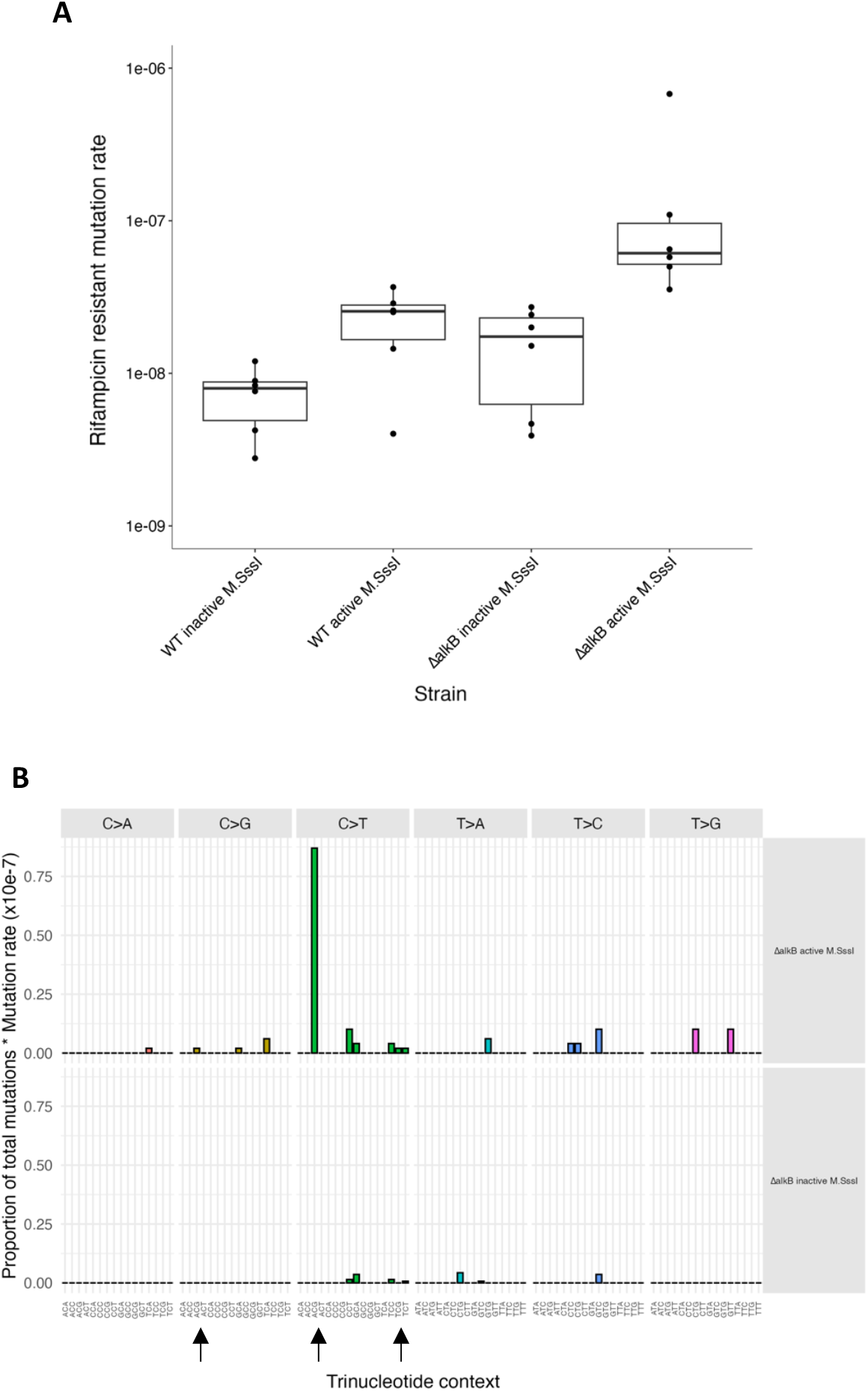
Constitutive M.SssI activity and loss of *alkB* increased mutation rate in Top10 *E. coli* but did not rescue the non-cytosine mutational signature. (A) WT and Δ*alkB* Top10 *E. coli* were transformed with active and inactive pM.SssI and plated on both rifampicin and LB media. The number of rifampicin-resistant mutant colonies was compared to total colonies on LB media that did not contain rifampicin to calculate a rifampicin-resistant mutation rate. Each individual point represents an independent biological replicate, each averaged over two technical replicates. (B) Mutational signatures in Δ*alkB* Top10 E. coli with active or inactive M.SssI. Signatures are categorised by substitution type and trinucleotide context, comprised of the 3’ and 5’ bases adjacent to the mutated base. This gives 96 possible signatures. The percentage of unique signature was multiplied by average mutation rate for each strain to give a qualitative contribution to overall mutation signature. Observed signatures occurring in the CpG context are indicated by black vertical arrows.

### RecA is not responsible for DNMT-induced AT mutations in *ΔalkB* **cells**

We compared the genotypes of Top10 and C2523 to identify potential sources of the differences in mutational spectra between them. Notably, C2523 contains WT RecA whereas Top10 carries the *recA1* point mutation commonly found in cloning strains. RecA is responsible for many aspects of DNA repair, most notably the SOS response, which controls the expression of DNA repair factors including translesion DNA polymerases that are responsible for mutagenesis (Maslowska *et al*, 2019).

To test whether lack of RecA might explain the absence of mutations at AT base pairs in Top10 *ΔalkB* cells, we introduced RecA on a plasmid (Badran & Liu, 2015) and evaluated the rifampicin resistant mutation rate. RecA transformation led to increased mutations in the presence of constitutive and inducible M.SssI, both in WT and *ΔalkB* cells (Figures 5A, B, 6A). However, there was no evidence of increased AT mutations in DNMT-expressing *ΔalkB* cells, either with inducible (Figure 5D) or constitutive M.SssI expression (Figure 6B). Thus, despite demonstrable effects of RecA expression in the Top10 cell background, this was not sufficient to induce DNMT-dependent mutagenesis at AT base pairs.

**Figure 5.**
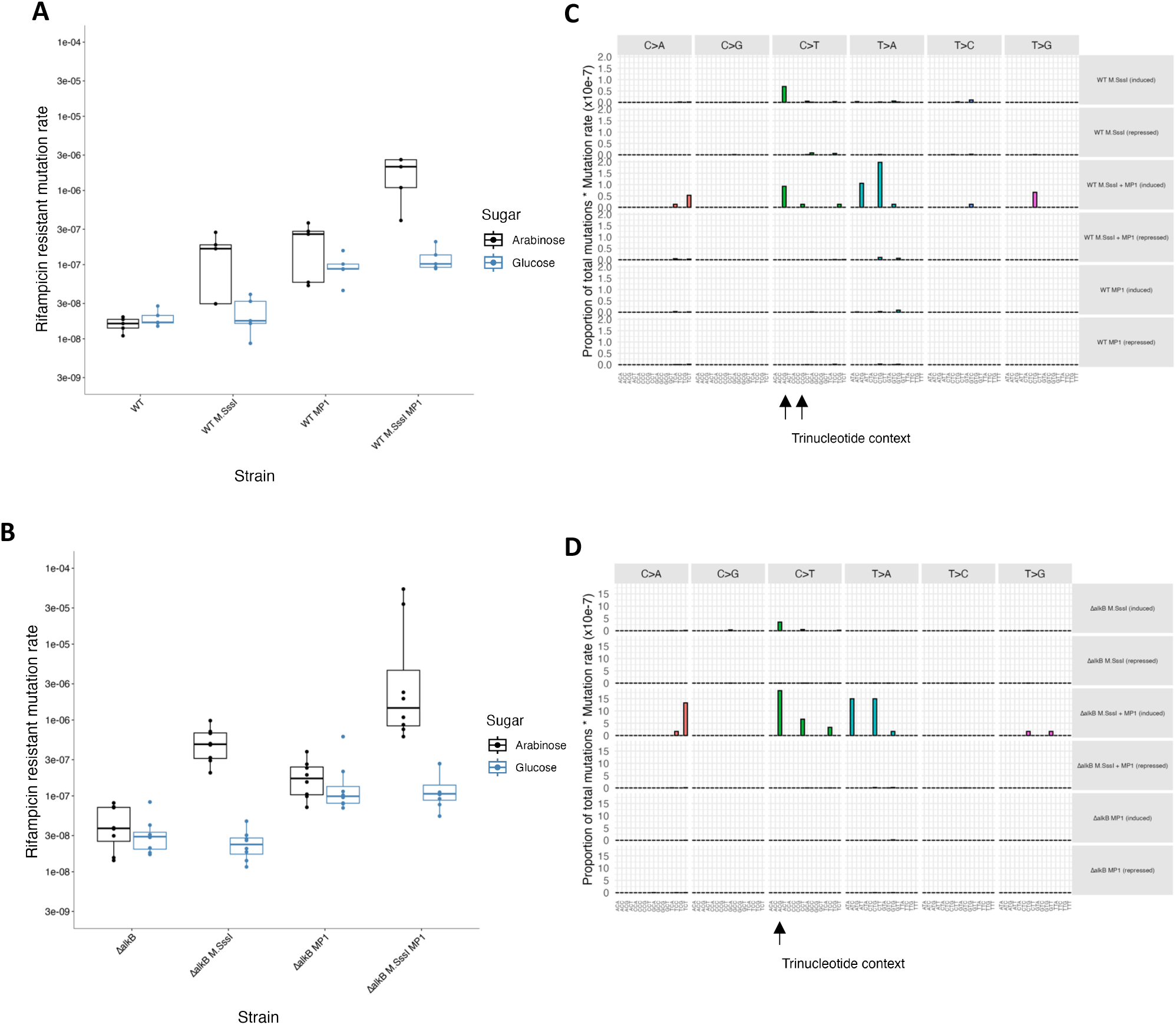
Addition of RecA in Top10 *E. coli* did not lead to induction of mutations at AT base pairs. Top10 *E. coli* were transformed with inducible pM.SssI and/or MP1. MP1 *encodes recA* and *umuC/D*, also under the control of an arabinose-inducible P_BAD_ promoter. Rifampicin resistant mutational frequencies were calculated for WT (A) and Δ*alkB* (B) cells. Mutational signatures for WT (C) and Δ*alkB* (D) Top10 *E. coli* transformed with inducible pM.SssI and/or MP1, treated with 0.5% arabinose (induced) or glucose (repressed). Signatures are categorised by substitution type and trinucleotide context, comprised of the 3’ and 5’ bases adjacent to the mutated base. This gives 96 possible signatures. The percentage of unique signature was multiplied by average mutation rate for each strain to give a qualitative contribution to overall mutation signature. Observed signatures occurring in the CpG context are indicated by black vertical arrows.

**Figure 6.**
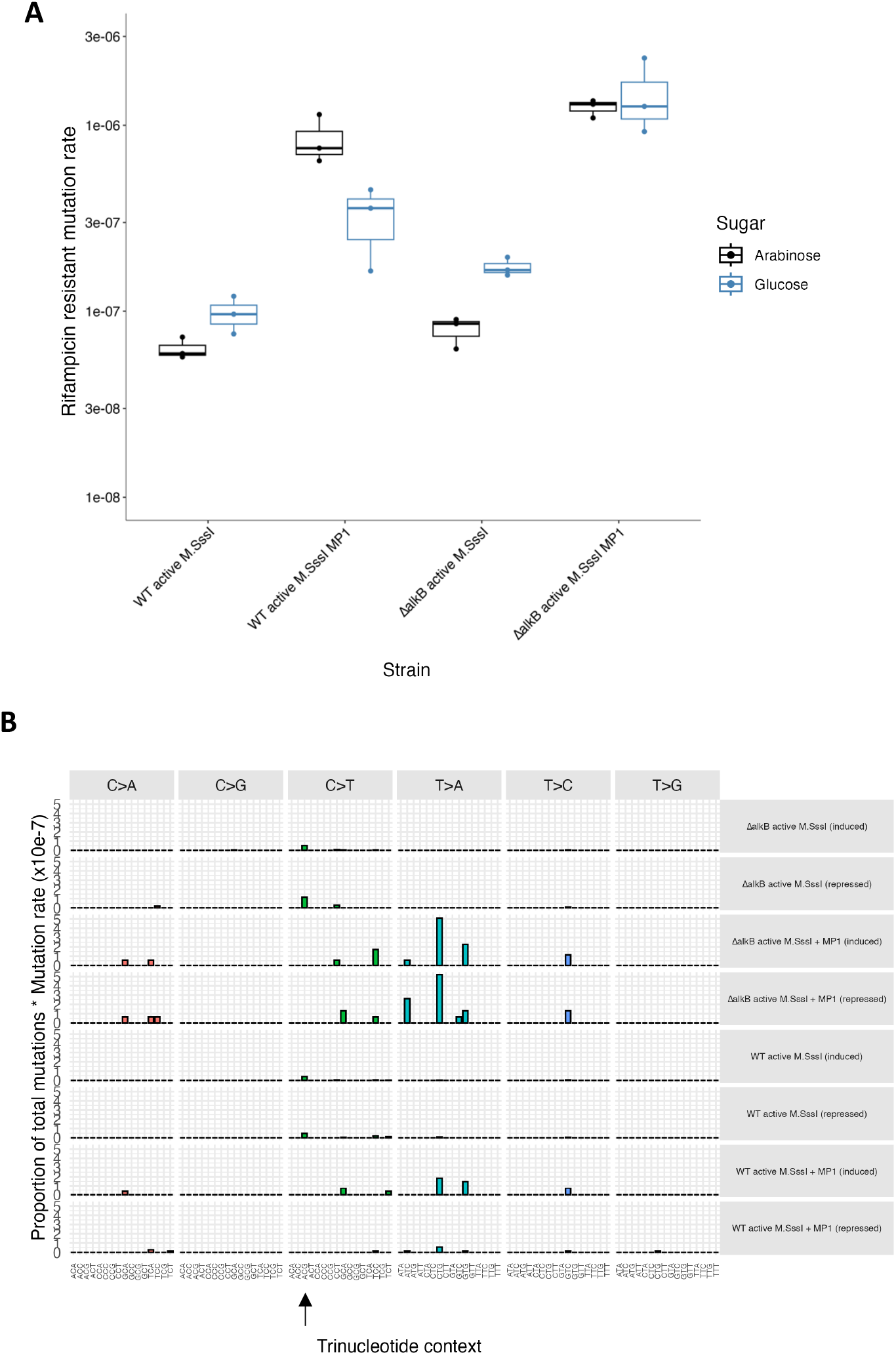
Introducing RecA into Top10 *E. coli* with constitutive M.SssI expression did not result in increased mutations at AT base pairs. (A) Rifampicin resistant mutation rates for WT and Δ*alkB* Top10 *E. coli* transformed with active pM.SssI +/-MP1. (B) Mutational signatures for WT and Δ*alkB* Top10 *E. coli* transformed with active pM.SssI +/-MP1. Separate signatures are shown for arabinose and glucose-treated conditions given that MP1 encodes *recA* under an arabinose-inducible P_BAD_ promoter. Signatures are categorised by substitution type and trinucleotide context, comprised of the 3’ and 5’ bases adjacent to the mutated base. This gives 96 possible signatures. The percentage of unique signature was multiplied by average mutation rate for each strain to give a qualitative contribution to overall mutation signature. Observed signatures occurring in the CpG context are indicated by black vertical arrows.

## Discussion

Epigenetic regulation through the induction of 5mC by DNA methyltransferases poses several distinct threats to genome stability. Here we explored the consequences of DNMT activity for mutagenesis. We showed that in WT cells, mutations at CG base pairs, specifically CG>TA, are induced by DNMT activity. However, we also identified an unexpected mutation pattern which appears only in the absence of the AlkB DNA repair enzyme. This mutation pattern involves mutations outside of CG sequences, instead forming AT->GC mutations. However, this effect is sensitive to cell background, and thus we cannot yet fully explain the molecular basis for these mutations.

### Mutagenesis due to DNMT activity

Here, we used a widely used assay to test for mutagenesis in *E. coli*, which involves selecting for resistant mutants to rifampicin which occur under spontaneous growth conditions. This assay is limited because not all possible mutations in *rpoB* lead to resistance to rifampicin so it does not provide accurate estimation of mutation rates or mutation spectra. However, we used the assay to compare mutation rates and spectra between different genotypes, which allowed us to draw qualitative conclusions about how mutagenesis is affected by DNMT expression.

We showed that mutations induced by DNMTs occur at CG base pairs and, unexpectedly, AT base pairs. Mutations at CG base pairs are consistent with the known activity of DNMTs. In WT *E. coli*, M.SssI introduced mutations at CG base pairs, within the context of CG dinucleotides. This suggests that the predominant mechanism for mutagenesis is deamination of methylated cytosine, which occurs at an elevated rate compared to unmethylated cytosine and is more challenging for cells to correct, as it produces thymine rather than uracil (Sarkies, 2022a). We observed an increase in this class of mutations in the *ΔalkB* background, particularly in the context of induction of DNMT activity in the Top10 cells. This suggests that AlkB prevents some CG>TA mutations from happening in WT Top10 cells. One explanation for this is the off-target activity of DNMTs in producing 3mC (Rošić *et al*, 2018). 3mC would normally be repaired by AlkB (Sedgwick, 2004), but in the absence of AlkB, it could cause mutations due to the fact that it blocks replication forks (Nieminuszczy *et al*, 2009). This might cause induction of translesion DNA synthesis, which could result in error prone replication (Sale *et al*, 2012). Erroneous incorporation of A opposite 3mC could lead to C>T mutations. TLS is known to bypass 3mC in vitro; however, it does not exclusively incorporate A (Furrer & Van Loon, 2014), so we might expect other mutations in addition to CG>TA, which we did not observe. This may be due to the limitations of the rifampicin resistance assay because of the limited number of mutations in *rpoB* that give rise to rifampicin resistance (Garibyan et al., 2003). In particular, the predominant CG>TA mutation in the CG context is a R->H substitution at the amino acid level. Further work incorporating unbiased mutagenesis assays such as generation of mutation accumulation lines could test this more thoroughly.

Mutations at AT base pairs differ from CG mutations in that they did not occur at all in the WT background, but only in backgrounds when AlkB is compromised. Additionally, we only observed these mutations in C2523 cells. What might their origin be? It seems unlikely that they result from alkylation damage induction by DNMTs: although 1mA is repaired by AlkB (Trewick *et al*, 2002), induction of 1mA by CpG specific DNMTs would require them to erroneously bind to A nucleotides, which seems implausible. Another possibility is that they are caused by translesion DNA polymerases extending bypass beyond 3mC sites induced by DNMT activity, in other words distal to CG dinucleotides. However, RecA, which is required for DNA damage-dependent induction of translesion DNA polymerases (Maslowska *et al*, 2019), does not seem to be responsible. At present, therefore, we cannot explain the molecular origin of these mutations. One possibility, however, is that failure to repair 3mC remaining in DNA in *ΔalkB* cells might lead to increased levels of single stranded DNA (Dinglay *et al*, 2000) which might increase the mutation rates of nucleotides close by. In this light it is interesting to note that mutation rates at nucleotides surrounding methylation sites are elevated in some species, although not in humans (Kusmartsev *et al*, 2020)

### Implications for the evolution of DNMTs

DNMTs are frequently lost across eukaryotic evolution. Here we show that there are mutagenic consequences associated with DNMT activity beyond what was already known from the direct effects of 5mC on deamination and miscoding. Most mutations are deleterious (Eyre-Walker & Keightley, 2007), so potentially the fitness costs associated with elevated mutagenesis might contribute to the loss of DNMTs, particularly in species where the eukaryotic homologue of AlkB that is specific for DNA (ALKB2 in eukaryotes) is also absent. It is worth noting that the extent to which mutation rate is under selection is debatable (Lynch *et al*, 2016), and likely depends strongly on the population size of the organism. The mutation rate is approximately 10-100 fold increased in our assay due to loss of *alkB* when DNMT is active. This might well be sufficient to overcome the drift barrier (Lynch *et al*, 2016), so that loss of DNMT in this background becomes selectively advantageous. It would be interesting therefore to examine mutation rates across eukaryotic species and link this to the presence or absence of DNMTs and ALKB2.

## METHODS

### 1.1. Bacterial strains

This study used the *E. coli* strains C2523 (New England Biolabs, Cat. No. C2523, genotype *fhuA2 [lon] ompT gal sulA11 R(mcr-73::miniTn10--TetS)2 [dcm] R(zgb-210::Tn10--TetS*) *endA1 Δ(mcrC-mrr)114::IS10*) and Top10 (ThermoFisher Scientific, Cat. No. C4040, genotype *F-mcrA Δ(mrr-hsdRMS-mcrBC) φ80lacZΔM15 ΔlacX74 recA1 araD139 Δ(ara-leu)7697 galU galK rpsL(StrR) endA1 nupG*). Both are *mcrA* deficient and therefore do not cleave methylated DNA. The study required a WT and Δ*alkB* version of each strain. Previously, P1 phage transduction was used to knock out (KO) *alkB* in C2523 and replace it with a kanamycin resistant cassette, using the Keio collection knock out strain as a donor (Baba *et al*, 2006). In some cases, an additional Δ*alkB* C2523 strain was used that possessed the gene KO but no KanR cassette. In Top10, *alkB* was knocked out via lambda red recombination, using a kanamycin resistance cassette from a donor plasmid, with flanking homology sequences to *alkB* (Datsenko & Wanner, 2000). Successful knockout of *alkB* was confirmed via PCR and selection on LB High Salt Agar + 50μg/mLl Kanamycin.

### 1.2. Plasmids

In order to introduce CpG methylation into *E. coli*, the bacterial CpG methyltransferase *m*.*sssI* from *Spiroplasma sp*. was cloned into pAIT2. This is a low copy number plasmid with both Kanamycin and Chloramphenicol resistance cassettes. In order to create a non-functional truncated control, the first 570bp of *m*.*sssI* were removed, which include the SAM binding domain, and cloned into pAIT2. Both the full length and truncated versions of *m*.*sssI* were under the control of a constitutive promoter. To allow for more precise control over induction of methylation, the full-length *m*.*sssI* was cloned into a pBAD plasmid with ampicillin resistance and the arabinose-inducible P_BAD_ promoter. This promoter is repressed in the presence of glucose, but activated upon arabinose addition, allowing for the induction of methyltransferase expression (Guzman *et al*, 1995). The inducible system was not functional in C2523, so was used solely in Top10 *E. coli*. Presence of methylation was determined by restriction digest with HpaII, which can only cleave unmethylated CCGG, and agarose gel electrophoresis. Top10 have a point mutation in *recA* (see genotype in 1.1). In order to introduce RecA into these cells, the high copy number plasmid pMP1 was used. Here, *recA* was under the control of the arabinose-inducible P_BAD_ promoter, with chloramphenicol resistance. pMP1 was a gift from David Liu (Addgene plasmid #69627 ; http://n2t.net/addgene:69627 ; RRID: Addgene_69627).

### 1.3. Bacterial media and culture conditions

All bacterial strains were cultured in Luria-Bertani (LB) media, with appropriate antibiotic selection (Kanamycin 50μg/ml, Chloramphenicol 30μg/ml, Ampicillin 100μg/ml) at 37°C and 220rpm. Strains were plated on LB High Salt Agar plates with antibiotics as above and incubated at 37°C overnight. WT and Δ*alkB* strains were made chemically competent with CaCl_2_ for transformation with the above plasmids (see 1.2). The transformed bacteria were plated on LB High Salt Agar with appropriate antibiotic selection and incubated overnight at 37°C. Additionally, in the case of *E. coli* containing the inducible methyltransferase and/or MP1, LB High Salt Agar and media were supplemented with 0.5% glucose to maintain promoter repression and suppress *m*.*sssI* and *recA* expression. Briefly, in order to induce expression, glucose-supplemented cultures were centrifuged and the pellet re-suspended in LB High Salt media + 0.5% arabinose.

### 1.4. Rifampicin resistance assay

In order to determine the difference in mutational burden caused by bacterial strains with and without methyltransferase activity, a rifampicin resistance assay was used. Rifampicin is an antibiotic that binds to the β subunit of RNA Polymerase. The gene *rpoB* encodes this subunit, and there are documented mutations in *rpoB* that cause resistance to Rifampicin (Garibyan *et al*, 2003). The mutational burden is approximated by comparing the number of rifampicin-resistant colonies to the total number of colonies on non-rifampicin media, giving a relative mutation rate.

WT and Δ*alkB* strains were transformed with either the full length or truncated form of *m*.*sssI*, or the inducible *m*.*sssI*. Where the introduction of RecA into Top10 was required, cells were transformed with both the inducible *m*.*sssI* and pMP1, and plated on LB High Salt Agar + antibiotics supplemented with 0.8% glucose. Individual colonies were inoculated overnight in LB High Salt media + antibiotics depending on plasmid content and strain background at 37°C, 220rpm. The next day, cultures were diluted 100x. For the constitutive system, cultures were allowed to grow to OD600 = 0.8 before centrifugation. For the inducible system and MP1, cultures were grown in 0.5% glucose to OD600 = 0.4. Then, cultures were centrifuged and the pellet re-suspended in LB High Salt media + 0.5% arabinose or glucose (control). All cultures were then allowed to grow to OD600 = 0.8-1.0 before centrifugation at room temperature. The pellet was then re-suspended in 300ul LB High Salt media. In order to measure the relative mutation rate for each strain, 100μl of undiluted culture was plated on 100μg/ml rifampicin plates and the remaining culture used to make ten-fold serial dilutions. An appropriate dilution was plated on LB High Salt Agar + antibiotic plates, to determine total cell count. The non-rifampicin plates were incubated overnight, while rifampicin plates were incubated for a total of 48 hours, both at 37°C. Relative mutation rates were calculated by dividing the number of mutant colonies by total number of colonies for each strain.

### 1.5. Colony PCR

Colony PCR was used following the rifampicin resistance assay to isolate genomic DNA and amplify a specific region, with the aim of identifying mutations via Sanger sequencing (see 2.7.). To liberate genomic DNA from inside the cell wall, individual rifampicin-resistant colonies were inoculated in dH_2_O and heated for 10 mins at 99°C. These were then centrifuged and the supernatant (DNA template) moved to a separate tube. PCR primers were used from Kurepina *et al*. that flanked a 510bp region of *rpoB*: rpoB_R_short: 5’ GTAGAGCGTGCGGTGAAAG 3’, rpoB_R_short: 5’ GCCTTTGCTACGGCAAGTTAC 3’ (Kurepina *et al*, 2022). PCR reactions were set up as follows: 1μl DNA template, 12.5μl 2x MasterMix (MedChemExpress, Cat No. HY-K0531), 1.25μl each of 10μM forward and reverse primers, 9μl dH_2_O. Cycles: 98°C 30 sec, then 25 x 98°C 10 sec, 68°C 30 sec, 72°C 30 sec, then 72°C for 2 minutes and 5°C for 10 minutes. PCR products were run on a 1.5% agarose gel to determine product size and specificity.

### 1.6. Sanger sequencing

Sanger sequencing was used to identify mutations arising in the fragments amplified by colony PCR (see 2.6.). 12μl of each PCR reaction (10ng/μl) was sent for clean-up and sequencing with Source Bioscience using the rpoB_R_short primer (3.2pmol/μl). Alignments were performed in Benchling using MAFTT v7 (Katoh & Standley, 2013).

## Supporting information

Mutation data collated

## DATA AVAILABILITY

Tabulated mutation data from analysis of Sanger sequencing is available as supplemental material

## ACKNOWLEDGEMENTS

This work was funded by the Royal Society (Investigating the mutagenic consequences of DNA alkylation damage by DNA methyltransferases; to PS) and the Wellcome Trust (How to Make a Parasite; to PS). We acknowledge Professor Nick Lakin, Professor Skirmantas Kriaucionis and Dr Joanna Krwawicz for advice on the project and members of the Sarkies group for helpful discussions.

## REFERENCES

Alexandrov LB, Kim J, Haradhvala NJ, Huang MN, Tian Ng AW, Wu Y, Boot A, Covington KR, Gordenin DA, Bergstrom EN, et al (2020) The repertoire of mutational signatures in human cancer. Nature 578: 94–101

Baba T, Ara T, Hasegawa M, Takai Y, Okumura Y, Baba M, Datsenko KA, Tomita M, Wanner BL & Mori H (2006) Construction of Escherichia coli K-12 in-frame, single-gene knockout mutants: the Keio collection. Molecular Systems Biology 2006 2:1 2: MSB4100050-

Badran, A. H., & Liu, D. R. (2015). Development of potent in vivo mutagenesis plasmids with broad mutational spectra. Nat Commun, 6, 8425. 10.1038/ncomms9425

Bewick AJ, Hofmeister BT, Powers RA, Mondo SJ, Grigoriev I V, James TY, Stajich JE & Schmitz RJ (2019a) Diversity of cytosine methylation across the fungal tree of life. Nat Ecol Evol 3: 479–490

Bewick AJ, Sanchez Z, Mckinney EC, Moore AJ, Moore PJ & Schmitz RJ (2019b) Dnmt1 is essential for egg production and embryo viability in the large milkweed bug, Oncopeltus fasciatus. Epigenetics Chromatin 12: 6

Bewick AJ, Vogel KJ, Moore AJ & Schmitz RJ (2017) Evolution of DNA methylation across insects. Mol Biol Evol 34: 654–665

Datsenko KA & Wanner BL (2000) One-step inactivation of chromosomal genes in Escherichia coli K-12 using PCR products. Proc Natl Acad Sci U S A 97: 6640–6645

Dinglay S, Trewick SC, Lindahl T & Sedgwick B (2000) Defective processing of methylated single-stranded DNA by E. coli alkB mutants. Genes Dev 14: 2097–2105

Dukatz M, Requena CE, Emperle M, Hajkova P, Sarkies P & Jeltsch A (2019) Mechanistic Insights into Cytosine-N3 Methylation by DNA Methyltransferase DNMT3A. J Mol Biol 431: 3139–3145

Eyre-Walker A & Keightley PD (2007) The distribution of fitness effects of new mutations. Nat Rev Genet 8: 610–618

Feng S, Cokus SJ, Zhang X, Chen P-Y, Bostick M, Goll MG, Hetzel J, Jain J, Strauss SH, Halpern ME, et al (2010) Conservation and divergence of methylation patterning in plants and animals. Proceedings of the National Academy of Sciences 107: 8689–8694

Furrer A & Van Loon B (2014) Handling the 3-methylcytosine lesion by six human DNA polymerases members of the B-, X- and Y-families. Nucleic Acids Res 42: 553–566

Garibyan L, Huang T, Kim M, Wolff E, Nguyen A, Nguyen T, Diep A, Hu K, Iverson A, Yang H, et al (2003) Use of the rpoB gene to determine the specificity of base substitution mutations on the Escherichia coli chromosome. DNA Repair (Amst) 2: 593–608

Guzman LM, Belin D, Carson MJ & Beckwith J (1995) Tight regulation, modulation, and high-level expression by vectors containing the arabinose PBAD promoter. J Bacteriol 177: 4121–4130

Holliday R, Kakutani T, Martienssen RA & Richards EJ (1987) The inheritance of epigenetic defects. Science 238: 163–170

Illingworth RS & Bird AP (2009) CpG islands - ‘A rough guide’. FEBS Lett 583: 1713–1720 doi:10.1016/j.febslet.2009.04.012 [PREPRINT]

Katoh K & Standley DM (2013) MAFFT Multiple Sequence Alignment Software Version 7: Improvements in Performance and Usability. Mol Biol Evol 30: 772–780

Krwawicz J, Sheeba CJ, Hains K, McMahon T, Zhang Y, Kriaucionis S & Sarkies P (2025) Introduction of cytosine-5 DNA methylation sensitizes cells to oxidative damage. Elife 13

Kurepina N, Chudaev M, Kreiswirth BN, Nikiforov V & Mustaev A (2022) Mutations compensating for the fitness cost of rifampicin resistance in Escherichia coli exert pleiotropic effect on RNA polymerase catalysis. Nucleic Acids Res 50: 5739–5756

Kusmartsev V, Drozdz M, Schuster-Böckler B & Warnecke T (2020) Cytosine Methylation Affects the Mutability of Neighboring Nucleotides in Germline and Soma. Genetics 214: 809–823

Law JA & Jacobsen SE (2010) Establishing, maintaining and modifying DNA methylation patterns in plants and animals. Nat Rev Genet 11: 204–220 doi:10.1038/nrg2719 [PREPRINT]

Lewis SH, Ross L, Bain SA, Pahita E, Smith SA, Cordaux R, Miska EA, Lenhard B, Jiggins FM & Sarkies P (2020) Widespread conservation and lineage-specific diversification of genome-wide DNA methylation patterns across arthropods. PLoS Genet 16

Liao J, Karnik R, Gu H, Ziller MJ, Clement K, Tsankov AM, Akopian V, Gifford CA, Donaghey J, Galonska C, et al (2015) Targeted disruption of DNMT1, DNMT3A and DNMT3B in human embryonic stem cells. Nature Genetics 2015 47:5 47: 469–478

Lindahl T (1996) The Croonian Lecture, 1996: endogenous damage to DNA. Philos Trans R Soc Lond B Biol Sci 351: 1529–1538

Liu P, Burdzy A & Sowers LC (2002) Substrate Recognition by a Family of Uracil-DNA Glycosylases: UNG, MUG, and TDG. Chem Res Toxicol 15: 1001–1009

Lynch M, Ackerman MS, Gout JF, Long H, Sung W, Thomas WK & Foster PL (2016) Genetic drift, selection and the evolution of the mutation rate. Nat Rev Genet 17: 704–714 doi:10.1038/nrg.2016.104 [PREPRINT]

Maslowska KH, Makiela-Dzbenska K & Fijalkowska IJ (2019) The SOS system: A complex and tightly regulated response to DNA damage. Environ Mol Mutagen 60: 368–384

de Mendoza A, Lister R & Bogdanovic O (2020) Evolution of DNA Methylome Diversity in Eukaryotes. J Mol Biol 432: 1687–1705 doi:10.1016/j.jmb.2019.11.003 [PREPRINT]

Nieminuszczy J, Mielecki D, Sikora A, Wrzesiński M, Chojnacka A, Krwawicz J, Janion C & Grzesiuk E (2009) Mutagenic potency of MMS-induced 1meA/3meC lesions in E. coli. Environ Mol Mutagen 50: 791–799

Okano M, Bell DW, Haber DA & Li E (1999) DNA Methyltransferases Dnmt3a and Dnmt3b Are Essential for De Novo Methylation and Mammalian Development. Cell 99: 247–257

Ponger L & Li WH (2005) Evolutionary diversification of DNA methyltransferases in eukaryotic genomes. Mol Biol Evol 22: 1119–1128

Rošić S, Amouroux R, Requena CE, Gomes A, Emperle M, Beltran T, Rane JK, Linnett S, Selkirk ME, Schiffer PH, et al (2018) Evolutionary analysis indicates that DNA alkylation damage is a byproduct of cytosine DNA methyltransferase activity. Nat Genet 50: 452–459

Sale JE, Lehmann AR & Woodgate R (2012) Y-family DNA polymerases and their role in tolerance of cellular DNA damage. Nat Rev Mol Cell Biol 13: 141–152 doi:10.1038/nrm3289 [PREPRINT]

Sarkies P (2022a) DNA Methyltransferases and DNA Damage. Adv Exp Med Biol 1389: 349–361

Sarkies P (2022b) Encyclopaedia of eukaryotic DNA methylation: from patterns to mechanisms and functions. Biochem Soc Trans 50: 1179–1190

Schulz NKE, Wagner CI, Ebeling J, Raddatz G, Diddens-de Buhr MF, Lyko F & Kurtz J (2018) Dnmt1 has an essential function despite the absence of CpG DNA methylation in the red flour beetle Tribolium castaneum. Sci Rep 8

Sedgwick B (2004) Repairing DNA-methylation damage. Nat Rev Mol Cell Biol 5: 148–157 doi:10.1038/nrm1312 [PREPRINT]

Sikora A, Mielecki D, Chojnacka A, Nieminuszczy J, Wrzesiński M & Grzesiuk E (2010) Lethal and mutagenic properties of MMS-generated DNA lesions in Escherichia coli cells deficient in BER and AlkB-directed DNA repair. Mutagenesis 25: 139–147

Tomkova M, McClellan M, Kriaucionis S & Schuster-Böckler B (2018) DNA Replication and associated repair pathways are involved in the mutagenesis of methylated cytosine. DNA Repair (Amst) 62: 1–7

Tomkova M, McClellan MJ, Crevel G, Shahid AM, Mozumdar N, Tomek J, Shepherd E, Cotterill S, Schuster-Böckler B & Kriaucionis S (2024) Human DNA polymerase ε is a source of C>T mutations at CpG dinucleotides. Nature Genetics 2024 56:11 56: 2506–2516

Trewick SC, Henshaw TF, Hausinger RP, Lindahl T & Sedgwick B (2002) Oxidative demethylation by Escherichia coli AlkB directly reverts DNA base damage. Nature 419: 174–178

Weber M, Hellmann I, Stadler MB, Ramos L, Pääbo S, Rebhan M & Schübeler D (2007) Distribution, silencing potential and evolutionary impact of promoter DNA methylation in the human genome. Nat Genet 39: 457–466

Zemach A, McDaniel IE, Silva P & Zilberman D (2010) Genome-wide evolutionary analysis of eukaryotic DNA methylation. Science (1979) 328: 916–919

